# A genome-scale antibiotic screen in *Serratia marcescens* identifies YdgH as a conserved modifier of cephalosporin and detergent susceptibility

**DOI:** 10.1101/2021.04.16.440252

**Authors:** Jacob E. Lazarus, Alyson R. Warr, Kathleen A. Westervelt, David C. Hooper, Matthew K. Waldor

## Abstract

*Serratia marcescens,* a member of the order Enterobacterales, is adept at colonizing healthcare environments and an important cause of invasive infections. Antibiotic resistance is a daunting problem in *S. marcescens* because in addition to plasmid-mediated mechanisms, most isolates have considerable intrinsic resistance to multiple antibiotic classes. To discover endogenous modifiers of antibiotic susceptibility in *S. marcescens,* a high-density transposon insertion library was subjected to sub-minimal inhibitory concentrations of two cephalosporins, cefoxitin and cefepime, as well as the fluoroquinolone ciprofloxacin. Comparisons of transposon insertion abundance before and after antibiotic exposure identified hundreds of potential modifiers of susceptibility to these agents. Using single gene deletions, we validated several candidate modifiers of cefoxitin susceptibility and chose *ydgH*, a gene of unknown function, for further characterization. In addition to cefoxitin, deletion of y*dgH* in *S. marcescens* resulted in decreased susceptibility to multiple 3^rd^ generation cephalosporins, and in contrast, to increased susceptibility to both cationic and anionic detergents. YdgH is highly conserved throughout the Enterobacterales, and we observed similar phenotypes in *Escherichia coli* O157:H7 and *Enterobacter cloacae* mutants. YdgH is predicted to localize to the periplasm and we speculate that it may be involved there in cell envelope homeostasis. Collectively, our findings provide insight into chromosomal mediators of antibiotic resistance in *S. marcescens* and will serve as a resource for further investigations of this important pathogen.

## Introduction

*Serratia marcescens*, a member of the order Enterobacterales, was historically regarded as an environmental bacterium with low inherent pathogenicity (1, 2). However, over the past 50 years it has been increasingly recognized as an important cause of invasive infections (3). *S. marcescens* can transiently colonize the gastrointestinal tract and skin, and is a frequent cause of sporadic healthcare-associated pneumonia, urinary tract and bloodstream infections (4–7). Because S*. marcescens* is commonly isolated from tap water (3) and clinical isolates frequently have high nucleotide identity to environmental isolates, it is thought that many infections result from sporadic exposures (8). However, since many isolates produce tenacious biofilms (9) and can have intrinsic resistance to common biocides (10, 11), it also causes hospital outbreaks (12), either through hand hygiene lapses or from a contaminated point source (13, 14). These outbreaks particularly affect vulnerable patients in adult and neonatal intensive care units (15, 16).

Antibiotic resistance (especially intrinsic resistance) is another crucial factor that allows *S. marcescens* to colonize hospital environments and infect vulnerable hosts. *S. marcescens* is intrinsically resistant to polymyxins (17, 18) and often has elevated minimal inhibitory concentrations (MICs) to tetracyclines, macrolides, nitrofurantoin, and fosfomycin (19). Importantly, *S. marcescens* also encodes a chromosomal Ambler class C beta-lactamase, AmpC, which at basal levels imparts resistance to penicillins and early generation cephalosporins (19). However, when treated with beta-lactams, *S. marcescens* clones with mutational derepression are selected, leading to overexpression of AmpC, which when expressed at high levels, can also impart resistance to late generation cephalosporins (20, 21). Disturbingly, there have also been increasing reports of dissemination of *S. marcescens* clones containing a mobilizable chromosomal genomic island (22) containing the class D beta-lactamase, SME, which efficiently hydrolyzes carbapenems (23–26). Infections with isolates that combine high-level expression of both AmpC and SME have been described (27); widespread dissemination of such highly beta-lactam resistant clones, on the background of fluoroquinolone non-susceptibility rates as high as 20% (https://sentry-mvp.jmilabs.com/) would leave clinicians to choose among only the most expensive and toxic, last-line treatment options.

To comprehensively identify loci that contribute both to basal growth and to antibiotic susceptibility in *S. marcescens,* here we use transposon insertion site sequencing (TIS) mutagenesis, a powerful approach that couples transposon mutagenesis with DNA sequencing. In TIS, a library of mutants is created where each bacterial cell harbors a transposon randomly inserted into the genome. Transposon insertions typically result in loss of function mutants, and in a high density library, genes for which transposon-insertion mutants are absent or underrepresented in the library are often critical for growth *in vitro* (sometimes termed “essential” genes). These genes may be potential antibiotic targets.

The library can additionally be subjected to biologically relevant conditions, and analyzing the abundance of mutants before and after exposure can identify genes that contribute to pathogen fitness in said condition. Mutants that are underrepresented after exposure to a condition of interest correspond to loci important for survival under that condition (29). This approach has facilitated rapid genome-scale identification of genes that contribute to phenotypes of interest (28), such as to identify genes that alter fitness in *ex vivo* and *in vivo* models of infection (30, 31), to investigate the function of uncharacterized genes (32), and to identify genes that alter antibiotic susceptibility. For example, TIS has identified novel modifiers of antibiotic susceptibility in many pathogens, including *Pseudomonas aeruginosa* (33), *Mycobacterium tuberculosis* (34), and *Klebsiella pneumoniae* (35).

Here, we created a dense transposon library in *S. marcescens* and used TIS to provide genome-scale insight into genes that contribute to *in vitro* growth as well as modifiers of cephalosporin and fluoroquinolone susceptibility. These analyses led to the identification of *ydgH*, a conserved gene that when deleted, leads to decreased cephalosporin susceptibility.

## Results

### Genes contributing to *in vitro* growth of *S. marcescens* ATCC 13880

We began by generating a high density transposon-insertion mutant library in a spontaneous streptomycin-resistant mutant of *S. marcescens* ATCC 13880, an environmental, non-clinical isolate that is the type strain for the species. Using a protocol adapted from *Escherichia coli* (36), we isolated nearly 2 million individual mutant colonies, each containing a genomic insertion of the mariner-based Himar1 transposon TnSC189 (37). Mariner transposons integrate at TA dinucleotides; the resulting pooled library included insertions at 57% of possible genomic TA sites. To ensure our library had sufficient complexity to allow subsequent analyses, we determined the percent of possible insertions achieved per gene. As expected for a high complexity library (28), a histogram of the resulting percentages revealed a bimodal distribution with a minor peak consisting of genes tolerating relatively few insertions and a major peak of disrupted genes centered around 70% TA site disruption (Figure 1A). Of the 4363 *S. marcescens* genes annotated, 4138 (94.8%) were isolated with at least one insertion. This allowed us to perform a comprehensive analysis of genes involved in *in vitro* growth of *S. marcescens*.

**Figure 1.**
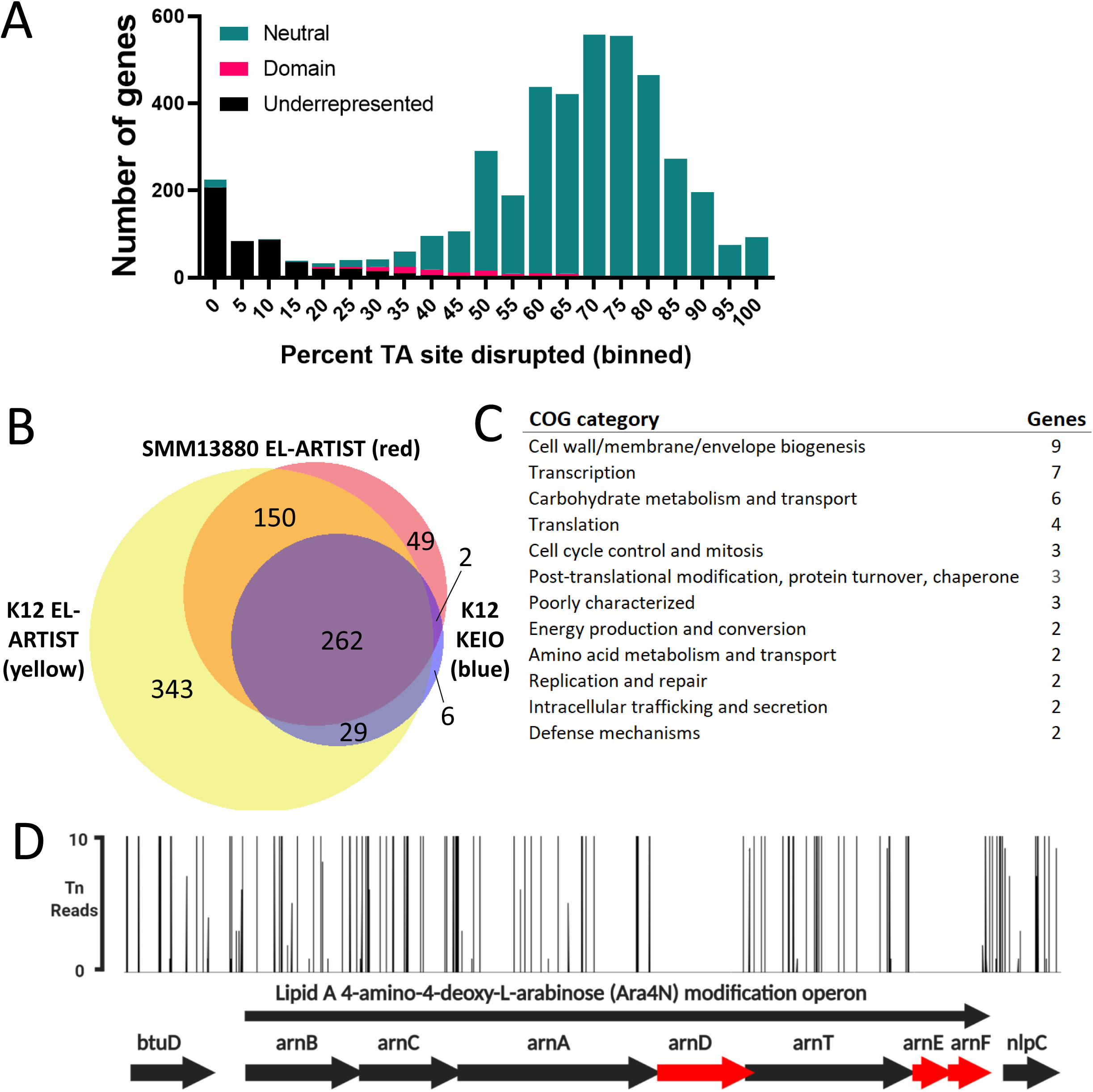
*S. marcescens* ATCC 13880 transposon-insertion sequencing reveals genes important for *in vitro* growth. A) *S. marcescens* genes binned by percent of TA sites within that gene disrupted. Overlayed is the hidden Markov model-based analysis assignment (“EL-ARTIST” pipeline) of whether insertions within the gene are underrepresented, vary by domain within the gene (“domain”), or are found to be distributed neutrally within the gene. The left histogram peak contains predominantly underrepresented genes where insertions are tolerated in only a low percentage of sites. B) Venn diagram illustrating overlap between *S. marcescens* ATCC 13880 (SMM13880) underrepresented genes (red) with *E. coli* K-12 underrepresented genes (yellow), with the K-12 KEIO collection as an additional comparator (blue). C) Genes underrepresented in *S. marcescens* ATCC 13880 but not in *E. coli* K12 either by EL-ARTIST or by the analysis resulting from the KEIO collection. COG categories of those genes are tabulated. D) Lipid A 4-amino-4-deoxy-L-arbinose modification operon, demonstrating that transposon insertions are underrepresented (red) in *arnD, arnE,* and *arnF*. *btuD* and *nlpC* are genes flanking this operon.

Using a previously developed pipeline that uses hidden Markov model-based analysis of insertions (38), we grouped genes into three categories: neutral, domain, and underrepresented. Neutral genes, such as *entE*, which synthesizes the siderophore enterobactin, may be crucial under certain physiologic conditions, but *in vitro*, tolerate transposon insertion throughout the span of the gene and are dispensable for growth (Supplemental Figure 1A, left). In contrast, underrepresented genes (often referred to as “essential” genes) such as *purA*, encoding the adenylosuccinate synthetase involved in purine metabolism, can sustain insertions at few or no sites while still allowing growth (Supplemental Figure 1A, middle). Finally, “domain” genes, such as *helD*, encoding a DNA helicase involved in unwinding of duplex DNA, can be found with insertions in certain domains or regions of the gene, but not others (Supplemental Figure 1A, right).

By this analysis, out of the 4363 *S. marcescens* genes annotated, we identified 483 underrepresented for growth in LB, and an additional 104 domain genes (Supplemental Table 1). The remaining 3776 genes were classified as neutral. Compared to a prior analysis in *S. marcescens*, we identified fewer genes as “essential” for *in vitro* growth, though a strict comparison is difficult since this prior effort used a clinical strain and utilized a low-density library with 32,000 unique insertion mutants (31, 39). Binning by cluster of orthologous gene (COG) functional category revealed that as expected, the most frequent categories for underrepresented genes were for core cellular processes such as translation (including many tRNA synthetases and ribosomal proteins), cell envelope biogenesis (including enzymes involved in peptidoglycan and lipopolysaccharide synthesis (LPS)), and coenzyme metabolism (including enzymes involved in central metabolism) (Supplemental Table 1). These categories are common for essential genes in other organisms (40).

*S. marcescens* was formerly classified as a member of the *Enterobacteriaceae*, but modern genome-based phylogenetics has re-assigned *Serratia* species into the sister *Yersiniaceae* family. We were eager to identify both underrepresented genes shared between *S. marcescens* and the common model *E. coli* lab strain, *E. coli* K-12, as well as those specific to *S. marcescens*, and so compared those identified here to those previously identified using the same approach in *E. coli* K-12 (36), as well as to those identified in *E. coli* K-12 using single gene knockouts (the “KEIO” collection described in (41)). Emphasizing the conserved physiology across the order Enterobacterales, of the 463 underrepresented genes we identified in *S. marcescens* that were also identified in *E. coli* (with an E value of 1*10^-10^ and percentage identity >30%), 412 (89%) (Figure 1B) were also underrepresented in *E. coli* K-12 by transposon insertion; of the 299 genes in *E. coli* K-12 identified as essential by single gene knockout that were also identified in *S. marcescens*, 264 (88%) were also underrepresented by our analysis (Figure 1B).

Genes underrepresented in *S. marcescens* (but not in *E. coli* K-12) (*n* = 49, Supplemental Table 2) were most commonly assigned to COG functional categories including cell envelope biogenesis (9 genes), transcription (7 genes), and carbohydrate metabolism and transport (6 genes) (Figure 1C). Additional investigation of these genes may identify divergent biology in *S. marcescens* that could be targets for novel narrow-spectrum antimicrobials. An intriguing example involves the lipid A 4-amino-4-deoxy-L-arabinose (Ara4N) modification (*arn*) operon. In *Enterobacteriaceae* as well as in *Pseudomonas aeruginosa,* this operon, when upregulated by PhoP/PhoQ, can lead to decreased susceptibility to cationic polypeptides like polymyxins by addition of positively charged Ara4n moieties to lipid A (42). In *S. marcescens,* which is known to be intrinsically resistant to polymyxins through *arn* (17), we detected an absence of insertions in *arnD*, *arnE*, and *arnF* (Figure 1D).

In addition to the *arn* operon described above, compared to *E. coli* K-12, we also found that two of the three components of the AcrAB-TolC RND family multidrug efflux pump were underrepresented in *S. marcescens* ATCC 13880*. mlaE,* which encodes the inner membrane permease component that facilitates transport of cell membrane lipids between the inner and outer membranes was underrepresented as well (Supplemental Table 2) (43). Components of peptidoglycan recycling are also underrepresented in *S. marcescens* (compared to *E. coli* K-12), including *nlpD*, which serves to activate cell wall amidases which act in daughter cell separation (44), and *murQ*, encoding a component of the cytoplasmic peptidoglycan recycling machinery (45).

### Antibiotic screen in *S. marcescens* ATCC 13880

We then subjected our high-density insertion library to antibiotics with the goal of discovering novel loci that modify antibiotic susceptibility. We focused our initial efforts herein on cephalosporins and fluoroquinolones as they are the antibiotic classes with fastest worldwide growth in consumption (46) and are currently clinically useful against serious *S. marcescens* infections (20, 47). Within the cephalosporins, we chose cefoxitin, an early generation cephalosporin that is readily hydrolyzed by the AmpC beta-lactamase and to which *S. marcescens* is relatively resistant, and cefepime, a late-generation cephalosporin that is negligibly hydrolyzed by AmpC and to which *S. marcescens* is relatively susceptible (21). We chose ciprofloxacin as among the fluoroquinolones, its MICs are typically lowest for *S. marcescens* (https://sentry-mvp.jmilabs.com/) (Figure 2A). We performed this screen at sub-MIC concentrations that over the course of the treatment resulted in ≤ 10-fold decrease in CFU (Figure 2B, Supplemental Figure 1B); this preserved library diversity so that when we sequenced the resulting libraries, we could identify enough TA insertion mutants per gene to allow us to identify genes that had small as well as large effects on growth and survival (28). Sequencing the resulting libraries allowed us to identify genes that when mutated by transposon insertion (presumably resulting in null or hypofunction) led to depletion or enrichment of that mutant when exposed to antibiotic. Genes with fewer insertion-mutants (depleted) represent candidate loci that support resistance, whereas genes with more abundant insertion-mutants (enriched) represent genes that may support susceptibility. As expected, under screen conditions, we measured robust induction of AmpC by cefoxitin (Supplemental Figure 1C) (21).

**Figure 2.**
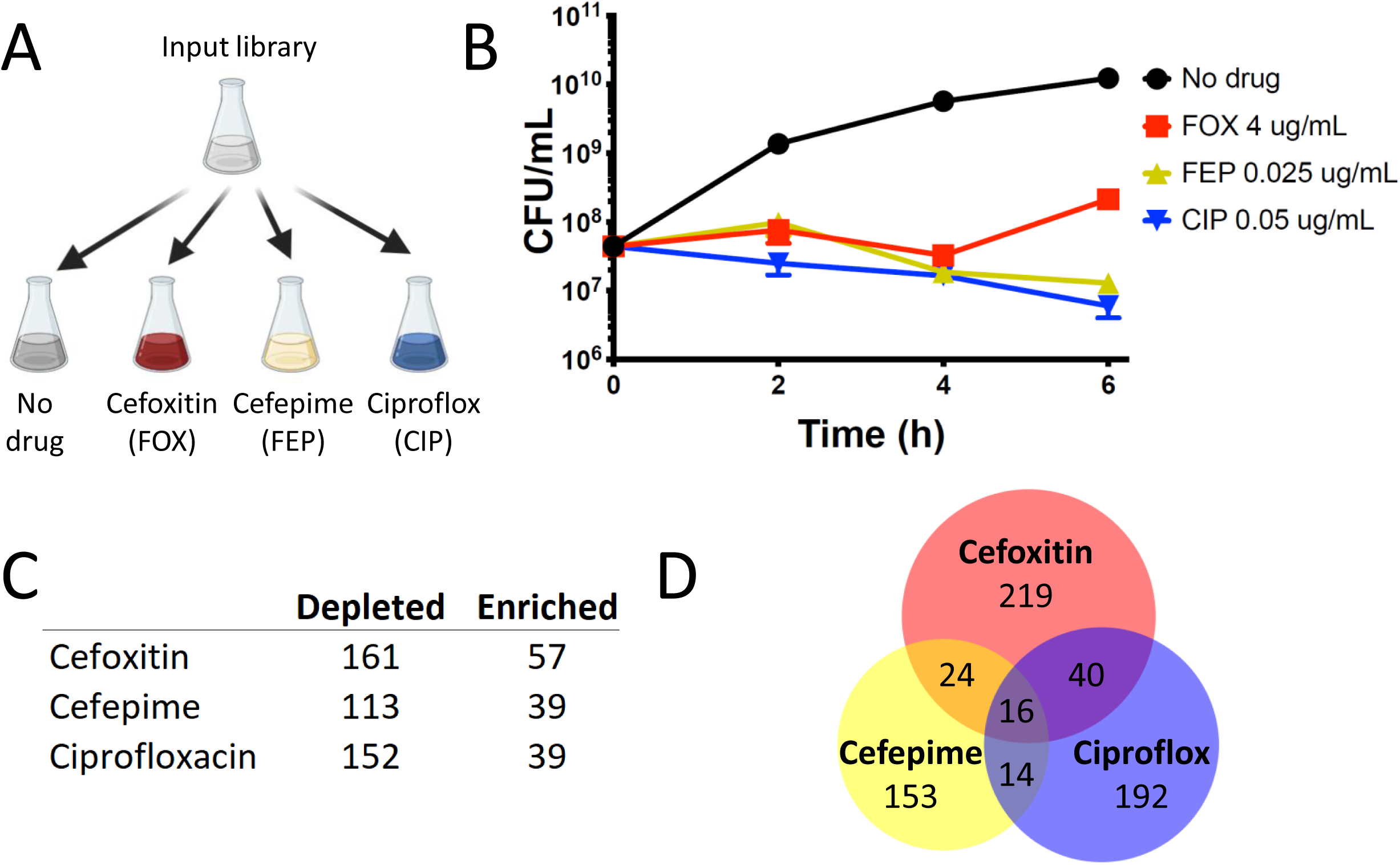
Antibiotic screen reveals *S. marcescens* genes important for growth and survival. A) Screen schematic: An input library (containing about 2 million insertion mutants) is grown to OD 0.1 and divided between 4 conditions for an additional 6 hours of growth: LB alone; LB + cefoxitin 4 ug/mL; LB + cefepime 0.025 ug/mL; and LB + ciprofloxacin 0.05 ug/mL. B) Under screen conditions, the library without drug selection undergoes more than 2 log expansion while those with antibiotic decrease in CFU to similar extent (about 0.5 log), with the library in cefoxitin undergoes expansion after 4 hours, likely due to upregulation of AmpC beta-lactamase. Libraries are harvested for analysis at 6 hours. C) Number of genes with insertion-mutants showing ≥ 4-fold change at 6 hours, compared to outgrowth in LB alone. D) Venn diagram illustrating genes showing coordinate enrichment or depletion.

Compared to outgrowth of the input library in no antibiotic, we found 57, 39, and 39 genes enriched and 161, 113, and 152 genes depleted in cefoxitin, cefepime, and the fluoroquinolone ciprofloxacin, respectively (with *p-*value ≤0.01 and ≥4-fold change in abundance) (Figure 2C, 2D). Many of the genes depleted in ciprofloxacin act in pathways related to the mechanism of action of fluoroquinolone antibiotics including those involved in DNA replication (such as *holE)* and DNA damage repair (such as *recD, recG*, and *xseA*), supporting the validity of our approach (Supplemental Figure 1D, Supplemental Table 3) (48, 49). Similarly, we identified many genes depleted in cefepime related to peptidoglycan homeostasis (such as *nlpD*, *pbpG*, *slt*, and *dacA*) and envelope integrity (such as *nlpA*) (Supplemental Figure 1E, Supplemental Table 3) (50).

As we would predict, in all 3 antibiotics we identified enrichment in insertion mutants in *ompF*, encoding an outer membrane porin that facilitates the permeation of cephalosporins and hydrophilic fluoroquinolones like ciprofloxacin (Supplemental Table 3) (51–53). All 3 also had enrichment of *lonP*, encoding a key bacterial serine protease whose deletion has been shown to lead to increased efflux of diverse antibiotics (54, 55). Interestingly, we also found enrichment in cefoxitin, cefepime, and ciprofloxacin in the setting of *slyA* insertion. SlyA is a member of the MarR family of transcriptional regulators known to be activated by small molecules; it is activated by salicylate and its best characterized role seems to be as a counter-silencer, alleviating H-NS-mediated repression (56). It has a diverse regulon (57), but has not previously been reported to be involved in regulation of antibiotic susceptibility. Further study is needed to characterize this and other genes we have identified (Figure 2D, Supplemental Table 3) in which insertion leads to coordinate depletion or enrichment in multiple antibiotics.

### Modifiers of cefoxitin susceptibility

To better understand modifiers of intrinsic resistance in *S. marcescens,* we focused our attention on modifiers of cefoxitin susceptibility, to which basal AmpC levels provide some baseline resistance. Of the 161 genes significantly depleted in cefoxitin (Figure 3A), we identified many predicted to participate in peptidoglycan homeostasis such as multiple membrane-bound lytic murein transglycosylases (*mltC*, *mltD*, and *mltF*), as well as penicillin-binding protein 1B (*mrcB*) and dihydrodipicolinate synthase (*dapA*) (Figure 3B). Importantly, mutants in *ampC* were also depleted, while we saw enrichment in mutants in the N-acetylmuramoyl-L-alanine amidase paralog that we have previously denoted as *amiD2* and suggested may be the most important for AmpC derepression in *S. marcescens* (58).

**Figure 3.**
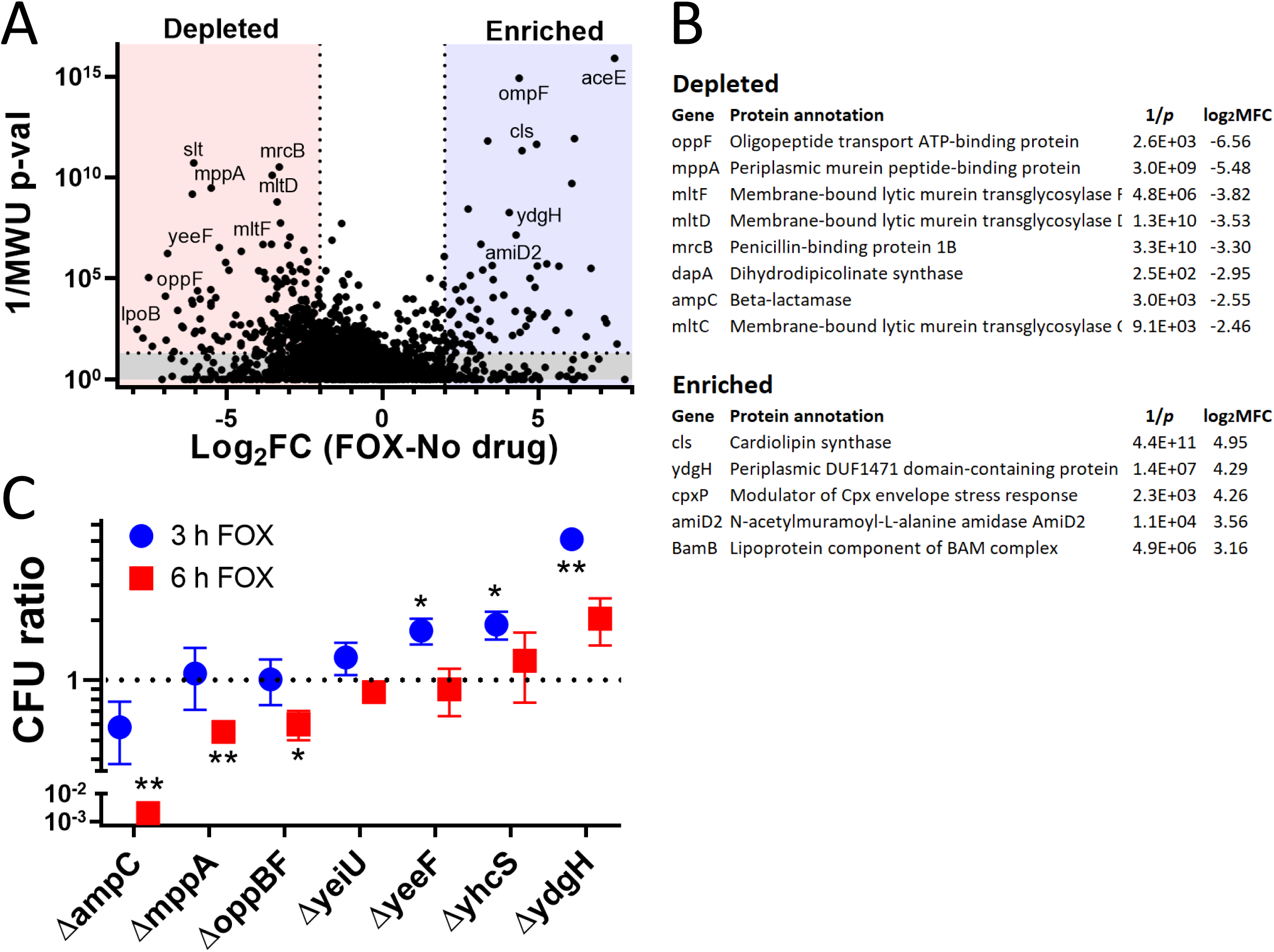
Whole genome screen for modifiers of cefoxitin susceptibility. A) “Volcano” plot illustrating candidate genes important for cefoxitin susceptibility. On the X-axis is the log_2_fold-change (Log_2_FC) in insertion-mutant abundance in cefoxitin (FOX) compared to no drug. The Y-axis is the inverse Mann-Whitney U *p-*value (1/MWU p-val), which roughly measures the concordance between mutants with insertions at individual TA sites across a gene. Genes were depleted (red) if Log_2_FC ≤ 2 and 1/MWU p-val ≥ 100. Genes were enriched (blue) if Log_2_FC ≥ 2 and 1/MWU p-val ≥ 100. B) Selected genes enriched or depleted in cefoxitin were tabulated. Genes important for envelope integrity and peptidoglycan recycling are depleted. *ydgH*, an enriched poorly-characterized gene, is also highlighted. C) Growth of various *S. marcescens* gene deletion mutants in cefoxitin 4 ug/mL. Growth is depicted as CFU_mutant_/CFU_Wt_ (in cefoxitin) / CFU_mutant_/CFU_Wt_ (in LB alone). Though no mutant had large defects in LB alone, due to stochastic errors in dilution to starting CFU, this improved the repeatability of the experiment.

Toward our aim of discovering novel modifiers of cefoxitin susceptibility, we specifically focused on genes in which insertion mutants were exclusively enriched or depleted in cefoxitin (and not also in cefepime and/or ciprofloxacin). We constructed in-frame deletions in 5 largely uncharacterized loci identified in our screen that were depleted (*mppA, oppBF, yeiU, yeeF,* and *yhcS*), and in one locus (*ydgH*), which was enriched. We compared the growth of the mutants in cefoxitin to that of wild-type *S. marcescens* ATCC 13880 in cefoxitin in a separate test tube. We used Δ*ampC* as a control. The final CFU ratio expressed as CFU_mutant_/CFU_wildtype_ is also corrected for any minor differences in growth that were observed in mutants in LB alone. CFU were determined after both 3 and 6 hours in cefoxitin (at the same concentration as used in the screen).

As expected, we observed a large progressive decrease in the CFU ratio in Δ*ampC* (Figure 3C). We also observed less pronounced but significant decreases in CFU ratios at 6 hours in Δ*mppA* and a mutant with a deletion of the entire *opp* operon, Δ*oppBF* (*oppB, oppC,* and *oppF* were all depleted in the screen). In contrast, deletions of *yeiU, yeeF*, and *yhcS,* in which insertions were depleted in our pooled screen, did not have reduced CFU ratios when these single gene deletions mutants were tested in isolation (Figure 3C).

### Δ*ydgH* has decreased susceptibility to cefoxitin

*ydgH*, which was enriched in the screen, had a large increase in CFU ratio at 3 hours of cefoxitin (Figure 3C). *ydgH* was originally identified through proteomic analyses as encoding a possible secreted effector in *Salmonella enterica;* however, YdgH was only detected in low abundance in samples from mutants deleted for a type III secretion system regulatory protein (and not in wild-type samples) (59). Efforts to identify cognate eukaryotic targets were unsuccessful (60).

In *S. marcescens* ATCC 13880, *ydgH* is predicted to encode a 951 amino acid protein. *ydgH* is upstream of the *ydgI* gene, which is predicted, by similarity to *P. aeruginosa* ArcD, to encode a putative arginine:ornithine antiporter (61). It is unlikely that *ydgH* and *ydgI* are part of an operon, as we identified high confidence σ^70^ promoters 5’ to both open reading frames as well as a transcriptional terminator 3’ to *ydgH* (Supplemental Figure 2) (62, 63). Furthermore, in contrast to *ydgH*, which contained enrichment of insertions throughout the gene in our TIS, there was no enrichment of insertions in *ydgI* (Figure 4A).

**Figure 4.**
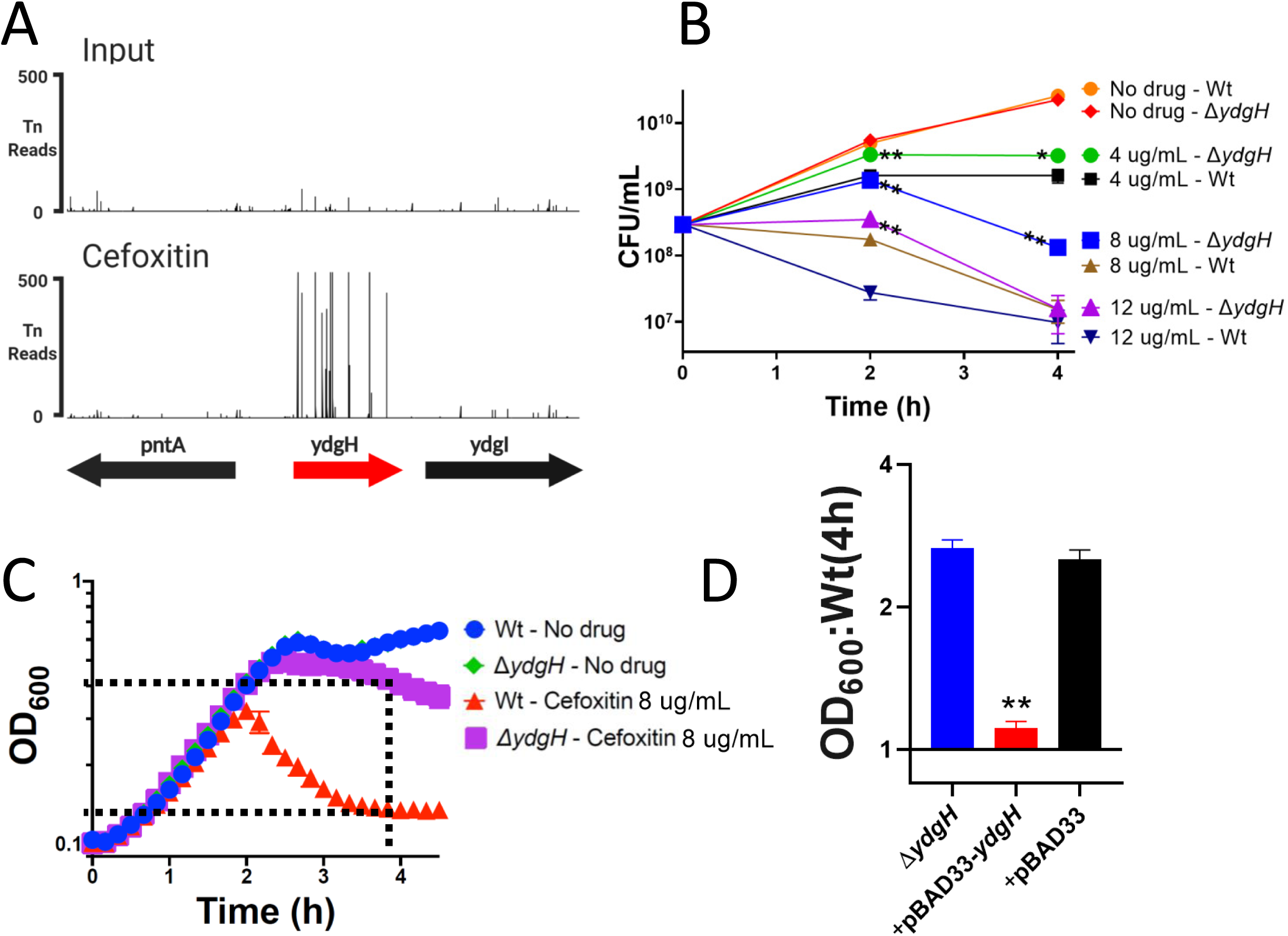
*YdgH* contributes to basal *S. marcescens* cefoxitin susceptibility. A) The *YdgH* locus on the Y-axis with transposon-insertion (Tn) reads on the Y axis, demonstrating the large enrichment of *YdgH* insertion mutants in cefoxitin (below, compared to the input library, above, on the same scale). B) Δ*ydgH* has increased growth in cefoxitin compared to Wt at multiple drug concentrations and at multiple time points. C) Schematic illustrating OD_6oo_ ratio used in subsequent figures: The OD_6oo_ of Δ*ydgH* at 4 hours is divided by that of Wt. D) Exogenous *ydgH* rescues the cefoxitin phenotype in Δ*ydgH*. Asterisks are * for *p* ≤ 0.05 and ** for *p* ≤ 0.01 by unpaired two-tailed *t* test.

We observed that Δ*ydgH* had significantly higher CFU than Wt across a range of cefoxitin concentrations at both 2 and 4 hours, but grew indistinguishably from Wt in the absence of cefoxitin (Figure 4B). We noticed smaller relative differences between Δ*ydgH* and Wt at later time points in these experiments, as well as those in Figure 3C, perhaps due either to compensatory upregulation of AmpC (at lower cefoxitin concentrations) or to delayed killing (at higher cefoxitin concentrations).

To enable higher-throughput screening of other compounds by spectrophotometry, for subsequent experiments, we used the OD_600_ ratio of Δ*ydgH* to Wt (as depicted in Figure 4C). We first used this approach to confirm that absence of *ydgH* was necessary and sufficient to confer enhanced growth in cefoxitin. Δ*ydgH* was transformed with either empty pBAD33 or pBAD33 containing the *ydgH* open reading frame; we observed that pBAD33-*ydgH* rescued the cefoxitin phenotype but the empty vector did not (Figure 4D).

### Δ*ydgH* has increased susceptibility to detergents

To identify if *ydgH* modifies susceptibility to other antibiotics, and to acquire initial clues to its function, we assayed the susceptibility of Δ*ydgH* to a range of antibiotics and detergents. For many compounds tested, there was a narrow concentration range that was sufficiently inhibitory to allow us to assay for potential effects. To enhance the accessibility of this large data set, we present the spectrophotometric ratio in the main text figures and the growth curves from which they are derived in the supplemental figures. The complete data at all concentrations tested are in Supplemental Table 4.

We began with beta-lactams; we found that in addition to the 2^nd^ generation cephalosporin cefoxitin, Δ*ydgH* also had significant reductions in susceptibility to the 3^rd^ generation cephalosporins moxalactam and ceftriaxone but not to the 1^st^ generation cephalosporin cephalexin or the anti-Pseudomonal cephalosporins cefepime or ceftazidime (Figure 5A, Supplemental Figure 3). There was not a prominent phenotype in the penicillins carbenicillin or piperacillin, or the carbapenems imipenem or meropenem (Figure 5A, Supplemental Figure 4). These distinct phenotypes are not attributable to differences in inherent susceptibilities to the assayed antibiotics, as the 3^rd^ generation cephalosporins, in which the mutant had phenotypes, and the anti-Pseudomonal cephalosporins, in which the mutant did not, were active across similar concentration ranges in *S. marcescens* ATCC 13880 (Supplemental Table 4). We hypothesized that the differences seen between the beta-lactams might be attributable to their being differential substrates for AmpC, but Δ*ydgH* did not have different AmpC activity compared to Wt (Supplemental Figure 4E).

**Figure 5.**
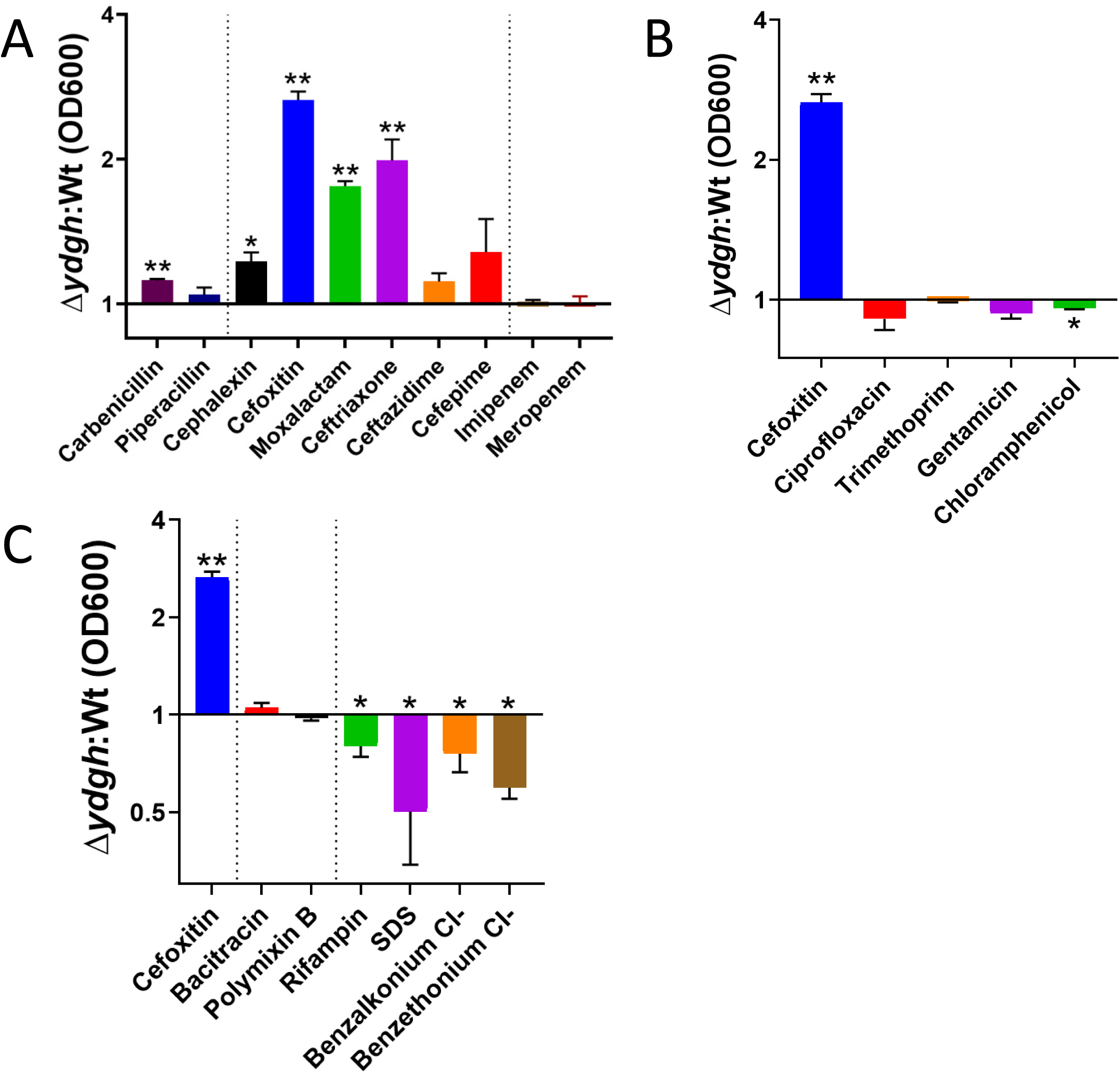
*YdgH* deletion leads to decreased cephalosporin susceptibility and increased detergent susceptibility. A) OD600 ratios demonstrating that Δ*ydgH* has decreased 2^nd^ and 3^rd^ generation cephalosporin susceptibility (including cefoxitin, moxalactam, and ceftriaxone) but no large differences in 1^st^ generation cephalosporins, anti-Pseudomonal cephalosporins, penicillins, or carbapenems. B) OD600 ratios demonstrating that Δ*ydgH* has no large differences in non-beta lactam antibiotics. Cefoxitin ratio reproduced for reference. C) OD600 ratios demonstrating that Δ*ydgH* has no large differences in bacitracin or polymyxin susceptibility, but has small but significant increases in susceptibility to rifampin, and more broadly to the detergents sodium dodecyl sulfate (SDS), benzethonium chloride and benzalkonium chloride. Cefoxitin ratio reproduced for reference. Asterisks are * for *p* ≤ 0.05 and ** for *p* ≤ 0.01 by unpaired two-tailed *t* test.

In contrast to the differential phenotypes observed with different beta-lactam antibiotics, we did not see a strong effect in antibiotics with cytoplasmic targets such as ciprofloxacin, trimethoprim, gentamicin, and chloramphenicol (Figure 5B, Supplemental Figure 5), nor to antimicrobials to which *S. marcescens* has high intrinsic resistance, such as bacitracin, and polymyxin B (Figure 5C, Supplemental Figure 6). Intriguingly, in contrast to the decreased susceptibility observed with 2^nd^ and 3^rd^ generation cephalosporins, we observed small, but significant *increased* susceptibility of the Δ*ydgH* mutant (compared to Wt) to rifampin, and more broadly to detergents including the anionic detergent sodium dodecyl sulfate (SDS), as well as the cationic detergents benzalkonium chloride and benzethonium chloride (Figure 5C, Supplemental Figure 6).

### Conservation within Enterobacterales

To gain further insight into this relatively uncharacterized gene, we performed a phylogenetic analysis and discovered that *ydgH* is closely conserved among the Enterobacterales (but not in other Gram-negatives) (Figure 6A). As expected, *S. marcescens* YdgH has greatest similarity to homologs identified in other *Yersiniaceae,* followed by the closely related sister families *Hafniaceae* and *Erwiniaceae,* followed by the more distantly related *Enterobacteriaceae*. We hypothesized that if the function of YdgH was conserved, we would observe similar phenotypes in distantly related Enterobacterales. To test this idea, we constructed in-frame deletions of *ydgH* in the pathogens *E. coli* O157:H7 EDL933 and *Enterobacter cloacae* ATCC 13047, and assayed the resulting mutants in ceftriaxone, to which *S. marcescens* Δ*ydgH* had decreased susceptibility, and in benzethonium chloride and SDS, to which *S. marcescens* Δ*ydgH* had increased susceptibility. We used ceftriaxone for these assays since *E. cloacae* has high-level intrinsic resistance to cefoxitin.

**Figure 6.**
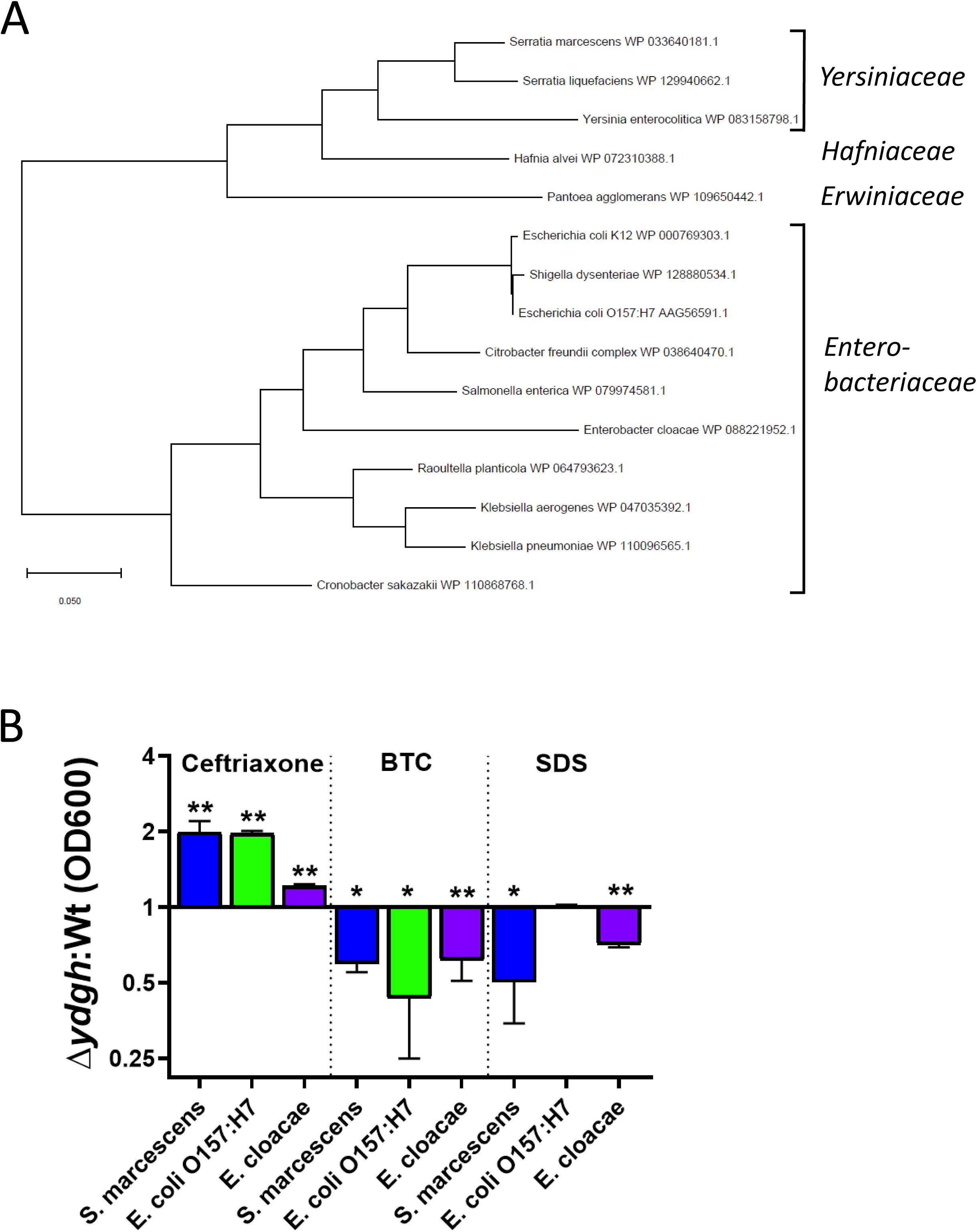
*YdgH* deletion results in conserved phenotypes in *Escherichia coli* O157:H7 EDL933 and *Enterobacter cloacae* ATCC 13047. A) The YdgH phylogeny was constructed based on amino acid substitutions by using the Maximum Likelihood method and JTT matrix-based model in MEGA-X. The relevant higher order families are indicated. B) All 3 mutants have decreased susceptibility to ceftriaxone (*E. cloacae* has high intrinsic resistance to cefoxitin so ceftriaxone was chosen) as well as increased susceptibility to benzethonium chloride. Growth of *E. cloacae* Δ*ydgH* was decreased in SDS though growth of *E. coli* O157:H7 was not. Due to inherent differences in Wt susceptibility, concentrations varied considerably between isolates and were ceftriaxone (4 ug/mL for *E. cloacae*, 0.04 ug/mL for *E. coli* O157:H7, 0.06 ug/mL for *S. marcescens)*, benzethonium chloride (BTC, 10 ug/mL for *E. cloacae,* 18 ug/mL for *E.* coli O157:H7, 52 ug/mL for *S. marcescens*), and SDS (0.8% for *E. cloacae,* 0.13% for *E. coli* O157:H7, 5% for *S. marcescens*).

We observed broadly consistent results, with *E. coli* O157:H7 Δ*ydgH* having similar phenotypes in ceftriaxone and benzethonium chloride (though not SDS), and *E. cloacae* Δ*ydgH* having similar phenotypes in benzethonium chloride and SDS (though with only a small difference in ceftriaxone) (Figure 6B).

## Discussion

Here, we present a genome-scale analysis of the essential genome of the type strain of *S. marcescens*, a medically important nosocomial pathogen. We report a curated resource of genes that alter susceptibility to beta-lactams and fluoroquinolones, arguably the two most useful antibiotic classes for treatment of *S. marcescens* infections. And, we validate and characterize *ydgH*, a largely uncharacterized gene conserved in the Enterobacterales, which when deleted leads to decreased susceptibility to 2^nd^ and 3^rd^ generation cephalosporins but increased susceptibility to ionic detergents.

A striking feature of the underrepresented genes we identified here in *S. marcescens* ATCC 13880 (compared to *E. coli* K-12) is their involvement in envelope biogenesis and homeostasis. These genes are involved in multiple structural and functional compartments including in LPS modification (*arnD, arnE,* and *arnF)*, phospholipid transport (*mlaE*), as well as peptidoglycan regulation (*murQ* and *nlpD*). These results hint at potentially important differences in *S. marcescens* and *E. coli* envelope physiology. It is known that in addition to the higher Ara4N content in Lipid A in *S. marcescens,* it also possesses additional core oligosaccharide substitutions (64, 65). We anticipate that further genetic investigations, including additional TIS to identify underrepresented genes in other relatively neglected Enterobacterales, as well as more detailed bioinformatic analyses using additional comparators, will suggest additional divergence in envelope biology, potentially enabling the development of narrow-spectrum antibiotics or antibiotic-adjuvants.

The high-density library we created should be a useful resource for investigation of other clinically-relevant phenotypes in *S. marcescens.* This approach has recently allowed identification of genes that facilitate bacteremia and the production of hemolysins in clinical strains of this pathogen (31, 66). Additional phenotypes crucial for pathogenesis, such as immune system evasion, biofilm formation, and colonization of target organs such as the bladder and lungs, will be of interest. The data set that we have generated here on cephalosporin- and fluoroquinolone-modifying genes should also serve as an important resource facilitating future investigations. Genome-scale antibiotic screens are an important addition to classical selection screens and sequencing of resistant clinical isolates because they can identify genes, such as *ydgH*, that regulates antibiotic susceptibility yet are not commonly seen clinically (perhaps due to conditional tradeoffs in fitness, like we identify here).

Our initial characterization of *ydgH* suggests that it does not act directly through the beta-lactamase AmpC; we show here that beta-lactamase activity is not increased in the Δ*ydgH* mutant. The increased susceptibility of Δ*ydgH* to detergents is likely an important clue regarding its function. YdgH is predicted to localize to the periplasm based on bioinformatic analysis and proteomics data sets (67, 68) and may in fact be present there in high concentrations in Enterobacterales (69, 70). One possibility is that in the periplasm, YdgH plays a role in the maturation of outer membrane-bound biomolecules. In support of this speculation, overexpression of *micA*, a small RNA that is a primary effector of the σ^E^ envelope stress response, leads to upregulation of *ydgH* (71). Alternatively, its absence may lead to envelope stress, leading to activation of σ^E^. Since downregulation of outer membrane porins that facilitate the entry of cefoxitin are a prime target of the σ^E^ response, this could explain both the increased susceptibility to detergents (and rifampin), and the decreased susceptibility to cefoxitin (and other cephalosporins particularly dependent on shared outer membrane porins for their entry).

Clearly, further mechanistic studies are needed to uncover the function of *ydgH,* as well as other genes that we identify here as modifiers of antibiotic susceptibility in *S. marcescens.* Such work will reveal novel bacterial cell biology as well as potentially tractable new antibiotic targets.

## Materials and Methods

### Transposon-insertion library construction

Conjugation was performed to transfer pSC189 (37) from *E. coli* SM10λpir to a spontaneous streptomycin-resistant mutant of *S. marcescens* ATCC 13880. Prior to conjugation, the two strains were cultured separately overnight in LB + chloramphenicol and LB + streptomycin, respectively, pelleted and washed twice in LB, resuspended 10-fold in LB, and combined 1:1 for final volume of 50 uL on a 0.45 um HA filter (Millipore) on an LB agar plate and incubated at 37°C for 1 hour. The filter was then eluted with 750 uL LB, and the filter washed twice more with 500 uL LB to ensure all transconjugants were eluted. 4 such reactions were then combined, and plated on 245×245 mm^2^ (Corning) LB-agar plates containing chloramphenicol and streptomycin. After 16 hours incubation at 37°C, colonies were harvested in LB + 25% glycerol (v/v) and stored at -80°C. This was repeated 3 times generating aliquots of “A,” “B,” and “C” libraries. The OD of the resulting libraries was measured (after appropriate dilution) prior to freezing and ranged from 56 to 108.

### Antibiotic screen

One aliquot each of the above libraries were thawed and combined (in proportion to its OD) in 50 mL of LB to yield a final OD of 3. 1 mL of above was then inoculated in 100 mL LB to roughly yield a culture with OD 0.03. The resulting culture was incubated with continuous agitation in 500 mL Erlenmeyer flask until OD of 0.1. 20 mL of the above culture was then added to 4-125 mL flasks, either an empty control flask or to flasks containing antibiotic to make a final concentration of cefoxitin 4 μg/mL, cefepime 0.025 μg/mL, or ciprofloxacin 0.05 μg/mL. Preparatory experiments had allowed estimation of the resulting CFU, so after 6 hours, to enable plating of roughly equal number of colonies, 100 μL of the LB culture (after dilution in LB), 5 mL of the cefoxitin culture (after washing and concentration in LB), and 20 mL of the cefepime and ciprofloxacin cultures (after washing and concentration in LB) were plated on LB agar without antibiotics. Surviving colonies were allowed to grow for 16 hours incubation at 37°C before colonies were harvested and stored as above. To determine beta-lactamase activity under screen conditions, samples from 2, 4, and 6 hours after incubation with antibiotic were taken, essentially as above except that a single aliquot of the “A” library was thawed rather than the 3 libraries combined. At the appropriate time points, a 1 mL aliquot was taken, pelleted at 13k RCF for 5 minutes at room temperature, resuspended in 1 mL 50 mM NaPO4 (pH 7) buffer, and flash frozen in liquid nitrogen and stored at -80 °C. After thawing, samples were lysed on ice using a Sonic Dismembranator with one pulse of 10 seconds on setting 8. Samples were clarified at 4°C 21k RCF for 10 minutes and the supernatants transferred to new tubes. Qbit protein assay (Invitrogen) was used to determine protein concentrations. 1 μg of total protein was added to 7.8 nmol nitrocefin (Sigma) and absorbance at 495 nm measured kinetically for 10 minutes at room temperature. The slope of the line from the first 5 data points was used to measure beta-lactamase activity (with 1 unit of activity representing hydrolysis of 1 μmole of nitrocefin per minute). Activity is expressed per gram of clarified lysate. AmpC is the only beta-lactamase identified in *S. marcescens* ATCC 13880 and Δ*ampC* mutants have essentially no beta-lactamase activity so beta-lactamase activity accurately represents AmpC activity. For the assay of AmpC activity in Δ*ydgH* to allow greater accuracy of a sample with lower relative activity, 2 μg of similarly obtained and processed mid-logarithmic samples were assayed.

### Characterization of transposon-insertion libraries

Libraries were prepared essentially as described in Warr et al (36). Prior to sequencing using a MiSeq V3 cartridge, equimolar DNA fragments for the original harvested library, and the resulting libraries after growth in LB alone, cefoxitin, cefepime, and ciprofloxacin were pooled after addition of barcodes to allow multiplexing. After trimming, reads were mapped, using Bowtie, allowing 1 mismatch, to the *S. marcescens* ATCC 13880 genome deposited below. Reads that mapped to multiple sites were randomly distributed between them. For analysis of underrepresented, domain, and neutral genes, EL-ARTIST was used after above. Artemis was used to generate TA insertion plots in Figures 1 and 4. Con-ARTIST was used to analyze conditional enrichment or depletion of transposon-insertion mutants, comparing the original input library to the library obtained after outgrowth in LB alone, cefoxitin, cefepime, and ciprofloxacin; these data are in Supplemental table 3. Genes conditionally depleted or enriched in antibiotics were compared to those conditionally depleted or enriched in the library outgrown in LB alone and if the original p-value was ≤0.01 and the adjusted fold-change ≥4, deemed significant. The “volcano” plots in Figure 3 and Supplemental Figure 1 are generated in this way. BioVenn was used to compare sets of conditionally enriched and depleted genes between antibiotic and between *S. marcescens* and *E. coli* K-12 data sets and to illustrate the results (72). Microsoft Excel was used to generate tables. Graphpad Prism was used to depict all remaining data.

### Molecular biology

Allelic exchange using pTOX3 was used to make all in-frame deletions (58) including *mppA* (WP_033640165.1), *oppB* (WP_004932150.1) *, oppC* (WP_004932149.1)*, oppD* (WP_016927496.1)*, oppF* (WP_004932143.1)*, yeiU* (WP_033639836.1)*, yeeF* (WP_004935642.1)*, yhcS* (WP_016929157.1)*, ydgH* (WP_033640181.1), and *ampC* (WP_033640466.1) all in *S. marcescens* ATCC 13880, as well as *ydgH* in *E. coli* O157:H7 EDL933 (WP_000769322.1) and *Enterobacter cloacae* ATCC 13047 (WP_013096870.1). pTOX3-derivative vectors were constructed essentially as described. Primer sequences are in Supplemental Table 5. Restriction enzyme-cut pTOX3 was incubated with purified AB and CD PCR products along with a half-reaction of HiFi DNA Assembly Master Mix (NEB) according to manufacturer’s directions. Reaction products were routinely desalted using “lily pad dialysis” (the entire above reaction volume placed on a slowly rotating 0.05 um HA filter (Millipore) floating on the surface of a MilliQ water) and subsequently electroporated directly into electrocompetent donor strains *E. coli* MFDλpir or SM10λpir. Selection and counter-selection were performed after (58). Deletion mutants were identified by colony PCR with subsequent Sanger sequencing of the reaction products. Primers used for construction of pBAD33-*ydgH* are in Supplemental Table 5. pBAD33-*ydgH* or pBAD33 were electroporated into Δ*ydgH* after (73). To ensure sufficient expression of YdgH, Δ*ydgH* pBAD33-*ydgH* (and empty vector) were induced in arabinose 1% (v/v) for 4 hours in mid-log phase prior to back-dilution and incubation with cefoxitin (or vehicle). To allow for confirmation of expression, a FLAG tag was cloned into the C-terminus. The YdgH phylogeny was constructed based on amino acid substitutions by using the Maximum Likelihood method and JTT matrix-based model in MEGA-X (74, 75).

### Δ*ydgH* chemical screen

Growth curves to identify phenotypes in Δ*ydgH* were performed using the indicated concentration of chemical (Sigma) in microplate format (Bioscreen C growth plate reader) using constant shaking setting at 37°C. Antibiotics were dissolved as recommended by the Clinical and Laboratory Standards Institute (CLSI) and stored in small aliquots at -80°C for up to 3 months.

### Data availability

*S. marcescens* ATCC 13880 was obtained from ATCC and a spontaneous streptomycin-resistant mutant isolated on LB + streptomycin 1000 ug/mL. This mutant grew indistinguishably from wild-type in the absence of streptomycin. DNA was isolated as directed using Qiagen DNeasy Plant Mini Kit and submitted to the University of Massachusetts Medical School Deep Sequencing Core which performed de novo assembly using PacBio sequencing and closed and finished using HGAP. The resulting genome, obtained from the core, was manually polished using Illumina reads and deposited (Accession number: CP072199). It is 99.93% identical to CP041233.1, which was not available at the at the time of sequencing. *S. marcescens* ATCC 13880 genes were identified using the BASys pipeline (incorporating Glimmer) and manually polished (76). The subsequent gene list was used to generate the look up table incorporated into EL-ARTIST and Con-ARTIST pipelines above. To ensure the accuracy of homology identification, for those *S. marcescens* identified as underrepresented, a translated protein BLAST was used to identify *E. coli* K-12 homologs and results manually adjudicated. COG categories for appropriate genes of interest were identified in the Database of Cluster of Orthologous Genes and manually tabulated (77). Reads for all sequenced libraries have been deposited in GEO (Accession number: GSE169651). UniProt and EcoCyc were frequently used to glean initial functional and sequence data about a particular gene product (78, 79). All statistics performed on processed data, depicted in figures were with an unpaired two-tailed *t* test. No correction for multiple comparisons were made. Significance asterisks are * for *p* ≤ 0.05 and ** for *p* ≤ 0.01. The *ydgH* gene schematic was constructed in Benchling using predicted sigma-70 promoter regions with a score > 90 in bacPP (62). Rho-independent terminators were identified using ARNold (63).

## Acknowledgements

We wish to thank members of the Waldor lab including Brandon Sit, Carole Kuehl and Troy Hubbard for helpful discussion. JEL has been supported by T32 AI-007061, the Harvard Catalyst Medical Research Investigator Training fellowship, and K08 AI-155830. MKW is supported by R01-AI-042347 and by the Howard Hughes Medical Institute.

## Supplemental figure legends

**Supplemental Figure 1 – A**) Depiction of represented neutral, underrepresented and domain genes. On the X-axis, the 3 genes are represented 5’ to 3’ to relative scale, with number of transposon (Tn) reads at that TA site on the Y axis. Many TA sites had more than 10 reads. B) OD_600_ of *S. marcescens* ATCC 13880 TIS library during the antibiotic screen. C) Beta-lactamase (AmpC) activity under conditions of the antibiotic screen. As expected, cefoxitin, which is a strong inducer of AmpC, results in large increases in beta-lactamase activity, as determined through hydrolysis of nitrocefin, a chromogenic cephalosporin substrate. D,E) “Volcano” plot illustrating candidate genes important for ciprofloxacin (in D) and cefepime (in E) susceptibility. On the X-axis is the log_2_fold-change (Log_2_FC) in insertion-mutant abundance in antibiotic (CIP and FEP) compared to no drug. The Y-axis is the inverse Mann-Whitney U *p-*value (1/MWU p-val), which roughly measures the concordance between mutants with insertions at individual TA sites across a gene. Genes were depleted (red) if Log_2_FC ≤ 2 and 1/MWU p-val ≥ 100. Genes were enriched (blue) if Log_2_FC ≥ 2 and 1/MWU p-val ≥ 100.

**Supplemental Figure 2 –** *YdgH* locus. Sigma-70 promoters with scores of 90 or greater in BacPP and rho-independent terminators identified by ARNold are indicated.

**Supplemental Figure 3 –** Growth curves of *S. marcescens* ATCC 13880 Wt or Δ*ydgH* in LB alone or in LB supplemented with the indicated concentrations (in ug/mL) of A) the 3^rd^ generation cephalosporin moxalactam; B) the 3^rd^ generation cephalosporin ceftriaxone; C) the 1^st^ generation cephalosporin cephalexin; D) the anti-Pseudomonal cephalosporins ceftazidime and E) cefepime. Informative concentrations used in calculating the OD600 ratios depicted in the main text are depicted. Results for the full range of concentrations tested are in Supplemental Table 4.

**Supplemental Figure 4** – Growth curves of *S. marcescens* ATCC 13880 Wt or Δ*ydgH* in LB alone or in LB supplemented with the indicated concentrations (in ug/mL) of the penicillins A) carbenicillin or B) piperacillin; and the carbapenems C) imipenem and D) meropenem. E) AmpC activity is not different in Δ*ydgH* compared to Wt, as measured by bulk nitrocefin hydrolysis of clarified supernatant.

**Supplemental Figure 5 -** Growth curves of *S. marcescens* ATCC 13880 Wt or Δ*ydgH* in LB alone or in LB supplemented with the indicated concentrations (in ug/mL) of the non-beta lactam antibiotics A) ciprofloxacin; B) trimethoprim; C) gentamicin and; D) chloramphenicol.

**Supplemental Figure 6 -** Growth curves of *S. marcescens* ATCC 13880 Wt or Δ*ydgH* in LB alone or in LB supplemented with the indicated concentrations (in ug/mL) of the antibiotics to which *S.* marcescens ATCC 13880 is intrinsically resistant, A) rifampin; B) bacitracin; and F) polymyxin B). Benzalkonium chloride and benzethonium chloride are depicted in D) and E). SDS concentrations in C) are (v/v).

**Supplemental table 1 –** Essential/underrepresented gene analysis of *S. marcescens* ATCC 13880 as determined through TIS and the EL-ARTIST pipeline. Summary statistics of essential/underrepresented genes underlying the analyses in Figure 1 are tabulated.

**Supplemental table 2 –** Essential/underrepresented genes unique to *S. marcescens* ATCC 13880 compared to both from *E. coli* K12 using either TIS and the EL-ARTIST pipeline or based on single gene deletion attempts resulting in the KEIO collection.

**Supplemental table 3** – TIS analysis identifying candidate genes important for outgrowth in no drug, cefoxitin, cefepime, or ciprofloxacin. Genes identified in all 3 antibiotics (compared to in no drug alone) are also tabulated.

**Supplemental table 4 –** Raw data from all final concentrations of chemical/antibiotic stressors used to generate main text figure 5 and the corresponding supplemental figures.

**Supplemental table 5 –** Primers used for creation of pTOX3 allelic exchange vectors and pBAD33-*ydgH*.

